# Non-antibiotic antimicrobial triclosan induces multiple antibiotic resistance through genetic mutation

**DOI:** 10.1101/267302

**Authors:** Ji Lu, Min Jin, Son Hoang Nguyen, Likai Mao, Jie Li, Lachlan J. M. Coin, Zhiguo Yuan, Jianhua Guo

## Abstract

Antibiotic resistance poses a major threat to public health. Overuse and misuse of antibiotics are generally recognised as the key factors contributing to antibiotic resistance. However, whether non-antibiotic, anti-microbial (NAAM) chemicals can directly induce antibiotic resistance is unclear. We aim to investigate whether the exposure to a NAAM chemical triclosan (TCS) has an impact on inducing antibiotic resistance on *Escherichia coli*. Here, we report that at a concentration of 0.2 mg/L TCS induces multi-drug resistance in wild-type *Escherichia coli* after 30-day TCS exposure. The oxidative stress induced by TCS caused genetic mutations in genes such as *fabI*, *frdD*, *marR*, *acrR* and *soxR*, and subsequent up-regulation of the transcription of genes encoding beta-lactamase and multi-drug efflux pump, together with down-regulation of genes related to membrane permeability. The findings advance our understanding of the potential role of NAAM chemicals in the dissemination of antibiotic resistance in microbes, and highlights the need for controlling biocide applications.

## 1. INTRODUCTION

The dissemination of antibiotic resistance has become a major threat to public health.^1^ Worldwide, each year about 700,000 people die from antimicrobial-resistant infections, and this mortality has been projected to reach 10 million per annum by 2050.^2^ The spread of antibiotic resistance has been attributed to the overuse and misuse of antibiotics in clinic settings, agriculture, and aquaculture.^3^

On a global scale, non-antibiotic, antimicrobial (NAAM) chemicals are used in much larger quantities than antibiotics, resulting in high residual levels of NAAM chemicals in the wider environment. For example, triclosan (TCS), a common biocidal agent used in over 2,000 kinds of products such as toothpaste and handwashing liquid,^4^ is widely detected in aquatic environments at *μ*g/L^5^ to mg/L levels,^6^ even up to 0.4 mg/L,^7^ Evidence suggests there are potential links between NAAM chemicals and antibiotic resistance.^8^ For instance, mupirocin-resistant^9^ and quinolone-resistant^10^ mutants were reported to exhibit decreased susceptibility to TCS, while TCS-resistant were found to have increased cross-resistance to ampicillin, ciprofloxacin^11^ and erythromycin.^12^ However, it remains unclear if NAAM chemicals such as TCS can directly induce antibiotic resistance. As a preventative policy, U.S. Food and Drug Administration (USFDA) has banned the addition of TCS to antibacterial soap.^13^ However, the lack of unequivocal evidence for NAAM chemicals inducing antibiotic resistance has prevented such a policy being adopted in other countries.

Here, we investigated the potential of TCS to cause antibiotic resistance. Wild-type *Escherichia coli* was exposed to TCS ranging from a sub-minimum inhibitory concentration (0.02 and 0.2 mg/L, which are environmentally relevant concentrations) to near lethal concentration (2 mg/L, Figure. 1A). We found that after 30 days treatment, 0.2 mg/L TCS increased the mutation frequency for multiple antibiotic resistance bacteria and the resistances were hereditable. Furthermore, we demonstrated that TCS at sub-MIC could induce mutation via increased Reactive Oxidative Species (ROS) generation and stress-induced mutagenesis (SIM). The subsequent mutations enhanced the expressions of beta-lactamase coding gene *ampC* and both local and global multidrug resistance regulator genes *acrAB*, *soxS* and *marAB* that initialized the translations of beta-lactamase AmpC and multidrug efflux pumps AcrAB-TolC. Together, TCS-induced mutations could induce resistance to antibiotics by increasing antibiotics efflux and antibiotic degradation.

**Figure 1.**
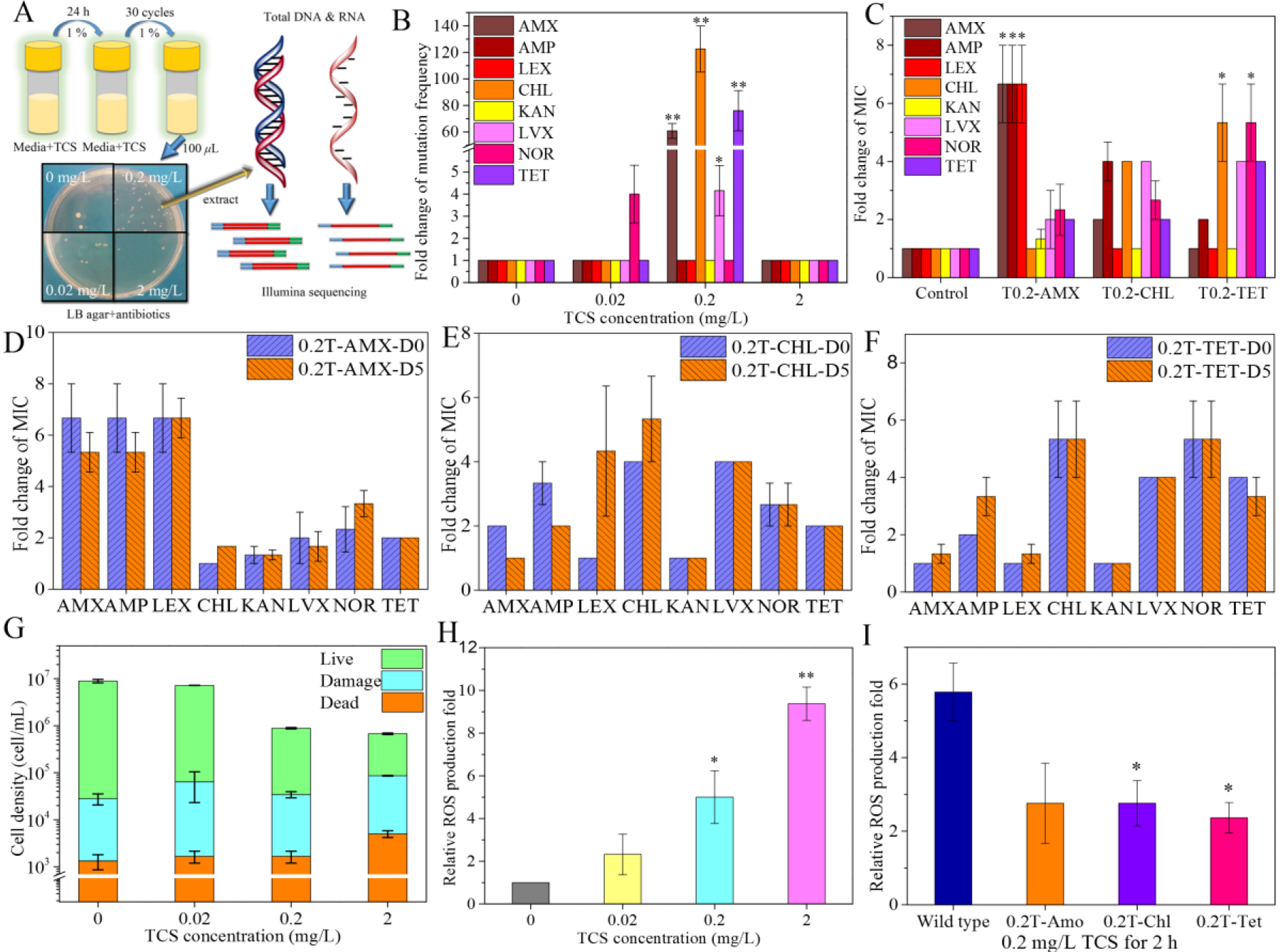
TCS caused reactive oxygen species (ROS) generation, and 30-day TCS treatment induced heritable multi-antibiotic resistance. (A) Set-up of experiment with TCS treatment. Every 24 h, 1% of TCS-treated *E. coli* culture was transferred into a fresh medium containing the same concentration of TCS. After 30 cycles, antibiotic-resistant strains were isolated by plating on LB agar supplemented with eight antibiotics, respectively (supplementary Table 1). Resistant strains isolated in this way were randomly selected to profile the MIC of multiple antibiotics (*n* = 8), followed by genome-wide DNA-and RNA sequencing to compare with wild-type *E. coli.* (B) Fold changes in mutation frequencies for amoxicillin (AMX), ampicillin (AMP), cephalexin (LEX), chloramphenicol (CHL), kanamycin (KAN), levofloxacin (LVX), norfloxacin (NOR) and tetracycline (TET) after 30-day exposure to TCS at different concentrations, relative to untreated *E. coli* (mean ± s.e.m., n ≥ 3), significance of difference with wild-type: \**p* < 0.05, ** *p* < 0.01 (Independent-samples *t*-test). (C) MIC fold increases for eight antibiotics determined for AMX-(0.2T-AMX), CHL-(0.2T-CHL) and TET-(0.2T-TET) resistant strains (mean ± s.e.m., n ≥ 5). All resistant strains showed statistically significant (\**p* < 0.05, ** *p* < 0.01, Independent-samples *t*-test) increases in MICs for multiple antibiotics. Fold change of MICs of 0.2T-AMX (D), 0.2T-CHL (E) and 0.2T-TET (F) strains at day 0 (D0) and after 5 days (D5) cultivation in the absence of antibiotics or TCS (mean ± s.e.m., n ≥ 3). The cultivation did not cause any significant changes (*p* > 0.05). (G) Concentration of viable cells (green), damaged cells (blue) and dead cells (orange) detected by flow cytometer after 2 h exposure to TCS (mean ± s.e.m., n = 3). (H) Relative ROS production by the wild-type *E. coli* after exposure to TCS for 2 h (mean ± s.e.m., n = 3), significance of difference with non-treated wild-type: \**p* < 0.05, ** *p* < 0.01 (Independent-samples *t*-test). (I) ROS production in wild-type *E. coli*, 0.2T-AMX, 0.2T-CHL and 0.2T-TET mutants following exposure to 0.2 mg/L of TCS for 2 h (mean ± s.e.m., n = 3), as enumerated by the ROS-positive percentage of fluorescence detected by flow cytometer (\**p* < 0.05, Independent-samples *t*-test).

## 2. MATERIALS AND METHODS

### Bacterial strains, triclosan, antibiotics

*E. coli* K-12 was purchased from American Type Culture Collection (ATCC 700926). Triclosan was purchased from Sigma-Aldrich (USA). Antibiotics: amoxicillin (AMX), cephalexin (LEX), tetracycline (TET), chloramphenicol (CHL), levofloxacin (LVX), and norfloxacin (NOR) were supplied by Sigma-Aldrich (USA). Kanamycin (KAN) was supplied by Astral Scientific (Australia), and ampicillin (AMP) was purchased from Gold Biotechnology (USA).

### Culture conditions, TCS exposure, and antibiotic-resistance determination

*E. coli* K-12 stock from −80 °C was cultivated on LB agar [lysogeny broth: 5 g/L yeast extract (Difco), 10 g/L NaCl and 10 g/L tryptone (Difco)] at 37 °C for 24h to isolate a single isogenic strain. The isolate was grown in liquid LB for 12 h at 37 °C to reach a bacterial cell concentration of 10^8^-10^9^ CFU/mL. For TCS exposure experiments, 50 *μ*L of the cell suspension was inoculated into 4.95 mL fresh liquid LB supplemented with different concentrations of TCS (0, 0.02, 0.2, and 2 mg/L, respectively) at 37 °C, shaken at 150 rpm, in triplicate. Every 24 h, 50 *μ*L of the cell mixture was transferred to another 5 mL tube containing 4.95 mL fresh, liquid LB with respective concentrations of TCS. This was repeated for 30 subculture cycles. At the end of the treatment, 100 *μ*L of each cell culture was plated on LB agar containing respective antibiotics at near MIC (Table S1) at 37 °C for 48 h, and then the number of colonies were counted. The colonies grown on the antibiotic-supplemented plates were considered to have resistance to the corresponding antibiotic. The mutation frequency was calculated by dividing the number of antibiotic-resistant colonies by the total bacterial count, which was enumerated from the LB agar without antibiotics.

### Determination of minimum inhibitory concentrations (MICs)

Following 30 days 0.2 mg/L TCS exposure, and subsequent cultivation on antibiotic selection plates (0.2T-AMX, 0.2T-CHL and 0.2T–TET), 5~8 antibiotic-resistant colonies were randomly picked, and incubated at 37 °C for 12 h in 2 mL of liquid LB. Using the selected strains, MICs were determined for eight antibiotics, respectively, including AMX, AMP, LEX, CHL, KAN, LVX, NOR and TET using an initial bacterial cell concentration of 10^6^ CFU/mL. Then, 15 *μ*L of this cell suspension was added to each well containing 135 *μ*L of serially, 2-fold diluted antibiotics in a fresh 96-well plate, followed by incubation at 37 °C for 24 h. The optical density (OD_600nm_) was measured using a plate reader Infinite^®^ 200 PRO (Tecan, Swiss). Each strain was tested in triplicate including sterilized PBS as a blank control. Fold changes in antibiotic-MICs were also calculated by dividing MICs of all treated mutants by the MIC of the wild-type *E. coli* (Figure 1C).

### Hereditary stability test

Antibiotic-resistant mutants (0.2T-AMX, 0.2T-CHL and 0.2T-TET) used for MIC profiling were cultivated in 5 mL liquid LB without antibiotics or TCS at 37 °C and 150 rpm. After 24 h, 1% of each liquid cell culture was transferred to fresh 5 mL treatment-free liquid LB and incubated under the same conditions. After five cycles, the MICs of eight antibiotics were tested respectively using the same method described previously, and the fold changes in MICs were determined. Each sample was tested in triplicate. MIC fold changes were determined for the cell cultures at day 0 and day 5 of incubation (Figure 1D, E and F).

### Live and dead cells percentages

The inhibitory effect of TCS on *E. coli* K-12 were investigated by staining with BacLight™ Bacterial Viability Kit (Invitrogen, USA). The LIVE/DEAD cell ratio of TCS-treated (0, 0.02, 0.2, and 2 mg/L TCS for 2 h) *E. coli* was then dual stained with propidium iodide (final concentration: 30 *μ*M) and SYTO 9 (final concentration: 5 *μ*M) in the dark at room temperature for 30 mins. The fluorescence was quantified by applying 500 *μ*L (10^6^ CFU/mL) of the stained samples to a CYTOFLEX flow cytometer (BD Biosciences, USA) with 488 nm excitation, and emissions were measured above 630 nm for red (PI) fluorescence and at 520 nm for green (SYTO 9) fluorescence. Untreated and heat-treated (2 h at 80 °C) cells were used as controls for intact and damaged cells, respectively.

### Detection of Reactive Oxygen Species (ROS)

To explore whether oxidative stress plays a role in promoting TCS-induced mutation, intracellular ROS formation was determined using the dye 2’,7’-dichlorofluorescein diacetate (DCF-DA, Abcam, UK), which can be oxidized by ROS into fluorescent compound, 2’, 7’– dichlorofluorescein (DCF), and measured with an Accuri C6 cytometer (BD Biosciences, USA). Briefly, bacterial cell suspensions (approximately 10^6^-10^7^ CFU/mL) were incubated with DCF-DA (at a final concentration of 20 *μ*M) for 30 min at 37 °C, shaken at 100 rpm in the dark. The bacterial cells were then directly treated with TCS (0.02, 0.2, and 2 mg/L) for 2 h at 37 °C, shaken at 100 rpm in the dark. A tert-butyl hydrogen peroxide (TBHP)-treated sample (50 *μ*M) was used as a positive control, and three non-TCS-treated samples as negative controls. The samples were then scanned by an Accuri C6 cytometer, and the DCF fluorescence (excitation at 488 nm/emission at 525 nm) was measured to deduce the ROS production level (Figure 1H). The relative ROS production level after dosage with 0.2 mg/L TCS for 2 h was determined for wild-type *E. coli* and 0.2T-AMX-, 0.2T-CHL-and 0.2T-TET-resistant strains which had been originally incubated with 0.2 mg/L TCS, 30 days (Figure 1I).

### DNA extraction, Illumina sequencing and data processing

MIC-profiled colonies from 0.2T–AMX-, 0.2T-CHL- and 0.2T–TET-resistant strains, and untreated *E. coli* K-12 were cultured in duplicate in 10 mL liquid LB at 37 °C for 16 h (shaken at 150 rpm) to reach 10^7^~10^8^ CFU/mL. Bacteria were then collected by 10 min centrifugation at 8000 × g, and genomic DNA was extracted using FastDNA™ SPIN Kit for Soil (MP, USA) following the manufacturer’s instructions. The NexteraXT DNA Sample preparation kit (Illumina, USA) was used to prepare a whole-genome shotgun library which was sequenced by AGRF (Brisbane, Australia) using a MiSeq instrument (Illumina, USA) with 150 bp paired-end sequencing, to a coverage of over 100-fold. A reference genome for *E. coli* strain K-12 was obtained from Genbank (Accession NC000913.3). The Illumina paired-end raw data was trimmed by Trimmomatic version 0.36^14^ to remove adapter and other illumine-specific sequences from the reads. After this, only properly paired reads were kept for further downstream analysis. These high read-depth data sets were aligned back to reference to study small variants e.g. Single nucleotide polymorphism (SNP), short indel (insertion or deletion), using BreSeq version 0.29.0.^15^ The allele frequency information was then extracted, and highly divergent loci were subjected to further analysis. Throughout the process, data visualization tools such as BRIG^16^ and Mauve^17^ were utilized for comprehensive comparisons and graphical plots.

### RNA extraction, genome-wide RNA sequencing and transcriptomic analysis

MIC-profiled 0.2T-AMX-, 0.2T-CHL-and 0.2T-TET-resistant mutants as well as the wild-type strain *E. coli* K-12 were cultured in triplicate in 10 mL liquid LB with 0.2 mg/L TCS at 37 °C for 8 h, shaken at 150 rpm. Bacterial cells were then collected by 10 min centrifugation at 8000 × g. Total RNA was extracted from the mutants using the QIAGEN miRNeasy Mini Kit (QIAGEN, Germany) manufacturer’s protocol with one extra bead-beating step to completely lyse the bacterial cells. RNA was treated with TURBO™ DNase (Ambion, USA) according to the manufacturer’s protocol, and integrity was confirmed via electrophoresis on a 2% agarose gel. Strand specific cDNA library construction and HiSeq 2500 (Illumina, USA) Illumina paired-end sequencing was conducted by Macrogen (Seoul, Korea). The NGS QC toolkit (version 2.3.3) was used to treat the raw sequence reads to trim the 3’-end residual primers and adaptors, following the removal of the ambiguous characters in the reads. Then, the sequence reads consisting of at least 85% bases were progressively trimmed at the 3’-ends until a quality value ≥ 20 were kept. Downstream analyses were performed using the generated clean reads of no shorter than 75 bp. The clean reads of each sequenced strains were aligned to the *E. coli* reference genome (NC_000913) using SeqAlto (version 0.5). The strand-specific coverage for each gene was calculated using Cufflinks (version 2.2.1), and to evaluate the differential t in triplicate bacterial cell cultures. CummeRbund package in R (http://compbio.mit.edu/cummeRbund/) were used to conduct the statistical analyses and visualization. Gene transcription was calculated as fragments per kilobase of a gene per million mapped reads (FPKM), a normalized value generated from the frequency of detection and the length of a given gene. Changes in expression values were calculated between wild-type *E. coli* control, 0.2 mg/L TCS-treated wild-type *E. coli*, and 0.2 mg/L TCS-treated 0.2T-AMX-, 0.2T-CHL-and 0.2T-TET mutants by determining the log2 fold change (LFC) of the averaged FPKM values of five, triplicate experiments. Genes with LFC of ≤-2 and ≥2, false discovery rate (FDR) of less than 0.05 and q value of less than 0.05, were labeled genes with highly differentially expression. Genes with LFC of <2 but ≥1, and ≤-2 but >-1 with q value of less than 0.05, and FDR of less than 0.05 were labeled moderately differentially expressed genes. Annotation of the differentially gene expressions was conducted by the online-curated Pathway Tools Genome Database, PseudoCyc (http://www.pseudomonas.com). The program Circos was applied to visualize the RPKM values for each gene amongst different samples.^18^

### Statistical Analysis

SPSS Statistics 24.0 (SPSS, Chicago, USA) was used for all data analysis. Significant differences were assessed using Independent-samples *t*-test. A value of *p* < 0.05 was considered significant, and a value of *p* < 0.01 was considered very significant.

### Data availability

All DNA sequencing data have been deposited with the National Center for Biotechnology Information (NCBI), and this is accessible through the SRA series (accession no. SRP124796). All RNA sequencing data have been deposited to the NCBI Gene Expression Omnibus, and this is accessible through the GEO series (accession no. GSE107048).

## 3. RESULTS AND DISCUSSION

### TCS treatment induces heritable multi-antibiotic resistance

We exposed wild-type *E. coli* to TCS with different concentrations (0, 0.02, 0.2 and 2 mg/L). Enumeration of antibiotic-resistant strains was carried out every 5 days by plating on LB agar supplemented with antibiotics (Table S1). The number of resistant colonies grown on plates containing different antibiotics was divided with total bacteria number to obtain the “mutation frequency”. The spontaneous mutation frequencies against eight types of antibiotics were established using the wide-type strain (non TCS-treated), which varied from 10^−7^ to 10^−8^ for different antibiotics. Then the mutation frequencies of TCS treated samples were compared with control and the increase of mutation frequency were calculated by the normalization to the spontaneous mutation frequency (Figure 1B). No significant increase in mutation frequency was detected until day 30, when 0.2 mg/L TCS significantly increased mutation frequencies against levofloxacin (LVX, 4.2 ± 1.1 times), amoxicillin (AMX, 60.8 ± 5.6 times), tetracycline (TET, 76.0 ± 15.2 times) and chloramphenicol (CHL, 122.6 ± 17.3 times), than that of the spontaneously resistant mutants without TCS treatment (\**p* < 0.05, \*\**p* < 0.01, Independent-samples *t*-test). In contrast, exposure to TCS at 0.02 and 2 mg/L did not lead to any significant increase in antibiotic-resistant mutants compared to the control without exposure to TCS (Figure 1B and Table S2).

To further confirm the development of antibiotic resistance, at least five colonies were randomly selected of each AMX- (labelled 0.2T-AMX), CHL- (labelled 0.2T-CHL) and TET- (labelled 0.2T-TET) resistant mutants induced by 30-day TCS treatment at 0.2 mg/L. The MICs of eight clinically important antibiotics were determined for the selected antibiotic-resistant mutants (Figure 1A). The 0.2T-AMX mutants expressed 6.7-fold increased MIC against the three beta-lactam antibiotics, ampicillin (AMP), cephalexin (LEX) and AMX. The 0.2T-CHL and 0.2T-TET mutants also showed increased MICs to multiple antibiotics of different categories (Figure 1C). Three types of resistant strains were scanned of antibiotic MICs on day 0 and day 5 cultivation (approximately 360 generations given a doubling time of 20 min) in the absence of TCS or antibiotics. All the antibiotic-resistant mutants on day 5 maintained similar antibiotic MICs compared to day 0 (Figure 1D-F), indicating genetically hereditary stability, and a possible mutagenic effect of TCS.

### TCS enhances generation of reactive oxygen species (ROS)

Previously, antibiotics were found to epigenetically generate antibiotic-resistant mutations via ROS stress response systems that damage DNA and the subsequent DNA repair system.^19^ Thus, we tested ROS generation and cell viability using flow cytometry, to verify whether TCS can induce oxidative stress in *E. coli*. We found that exposure to TCS at 0.2 and 2 mg/L for 2 h induced inhibitory effects on *E. coli* growth, the total cell concentration reduced from 8.86 × 10^6^ ± 7.79 × 105 cell/mL (0 mg/L TCS) to 8.17 × 10^5^ ± 3.03 × 10^4^ cell/mL (0.2 mg/L TCS) and 5.06 × 10^5^ ± 2.60 × 10^4^ cell/mL (2 mg TCS) (Figure 1G). Meanwhile, TCS exposure increased dead cell number from 1.33 × 103 ± 4.71 × 10^2^ cell/mL (0 mg/L TCS) to 1.66 × 10^3^ ± 4.71 × 10^2^ cell/mL (0.2 mg/L TCS) and 5.00 × 10^3^ ± 8.16 × 10^2^ cell/mL (2 mg TCS). More importantly, both 0.2 and 2 mg/L TCS treatments significantly increased generation of ROS for 5.00 ± 1.24 folds and 9.37 ± 0.78 folds, respectively (Figure 1H). In contrast, TCS treatment at 0.02 mg/L did not trigger significant ROS generation (Figure 1H) or cell inhibition (Figure 1G), which is consistent with a previous report(10) that TCS at 0.03 mg/L did not induce antibiotic resistance. Noticeably, although TCS dosage of 2 mg/L increased the ROS production, it did not increase mutation frequency (Figure 1B), likely because this near-MIC dosage of TCS reduced the frequency of accumulating mutations by decreasing the cell proliferation rate. These results indicate that there might be a threshold TCS concentration for the generation of antibiotic resistance. Moreover, TCS-induced mutants generated less ROS compared to wild-type *E. coli* (Figure 1I), suggesting mutants have developed resistance to TCS-induced oxidative stress.

### TCS triggers genetic mutations

To reveal the genetic changes involved in TCS-induced antibiotic resistance, we conducted genome-wide DNA sequencing on three types (0.2T-AMX, 0.2T-CHL and 0.2T-TET) of replicated TCS-induced mutants (*n* = 6) and wild-type *E. coli* (*n* = 2) (Figure 2). Sequencing of 6 antibiotic-resistant mutants generated after exposure to TCS at 0.2 mg/L revealed 14 genetic changes in 11 genes and 9 mutations in intergenic spacers compared with the untreated *E. coli* (Figure 2 and Tables S3). All sequenced mutants displayed the +A insertion in the *insB*−1 gene as well as substitution mutations in the *fabI* gene. TCS inhibits bacterial fatty acid synthesis by binding enoyl reductase FabI which encoded by *fabI* gene,^20^ therefore missense mutation in the *fabI* gene could reduce TCS efficacy by altering the structure of the target FabI protein.^21^ In addition to these shared mutations, some strain-specific mutations were also identified. For example, the substitution mutations of *acrR* (L65R), *citC* (A346T) and *soxR* (R20S) genes were only identified in the 0.2T-CHL strains. For the 0.2T-TET strain, a Δ1 bp frameshift in *insl*−1 gene and a substitution mutation in *marR* gene (T72P) were unique compared to other strains. Lastly, the 0.2T-AMX strain were detected with genetic changes include the substitution mutations in *aaeB* (V311G) and *rpoD* genes (D445G), an insertion in *frdD* gene (345/360 nt) as well as an insertion in intergenic spacer between *rrlG* and *rrfG* genes (Figure 2C).

**Figure 2.**
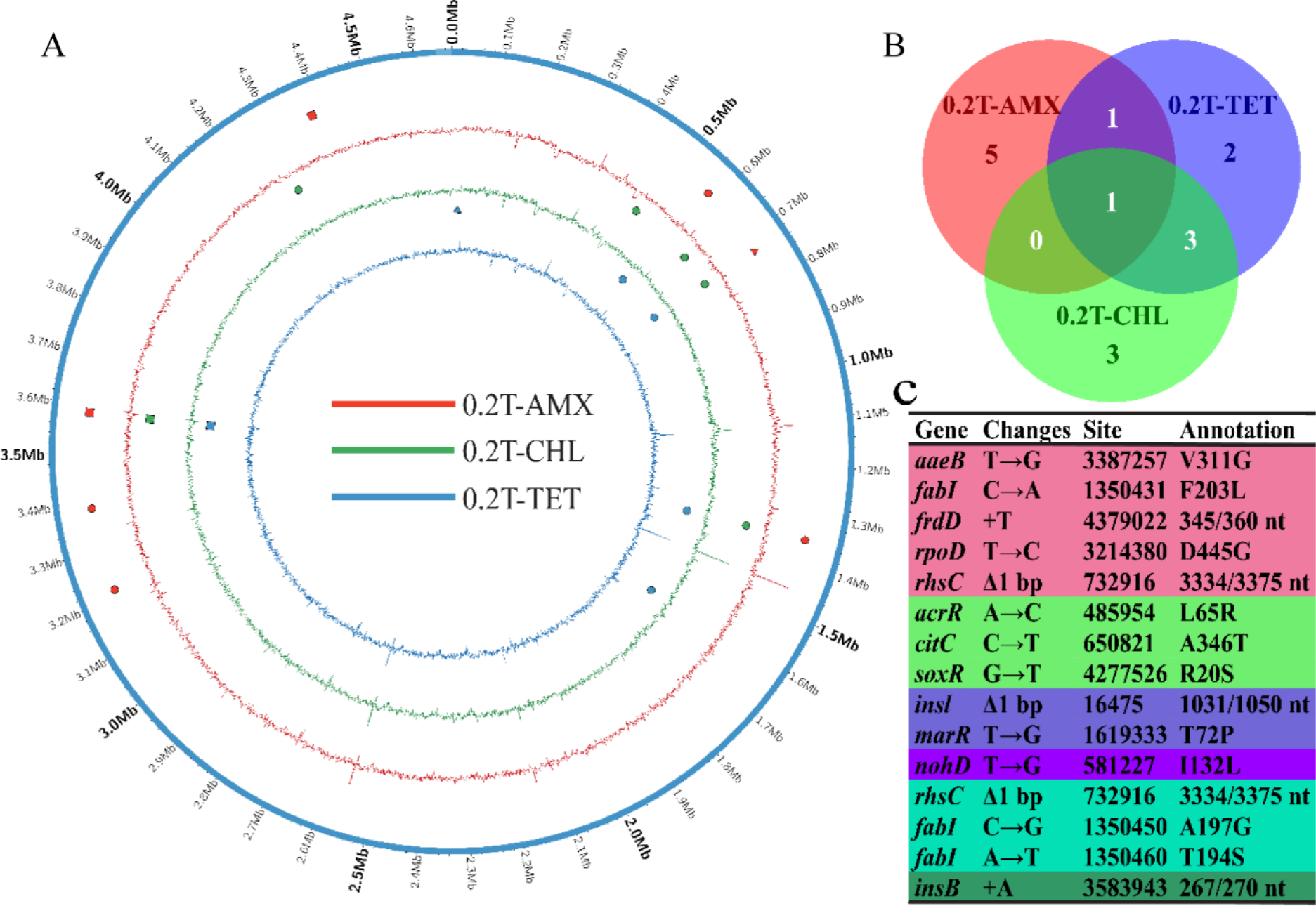
TCS-induced genetic mutations. (A) Genetic changes identified by Illumina whole genomic analysis in three types of mutants (*n* = 6, from inside to outside: 0.2T-AMX, 0.2T-CHL and 0.2T-TET) selected with exposure to 0.2 mg/L TCS for 30 days, compared with wild-type *E. coli* (*n* = 2). The outer circle represents the 4.63 Mb *E. coli* reference genome. ⚫ represents single nucleotide polymorphism, ■ represents a nucleotide insertion and ▲ represents a nucleotide deletion at the corresponding sites of the genome. (B) Venn diagram showing the number of gene mutations in 0.2T-AMX (red), 0.2T-CHL (green) and 0.2T-TET (blue) mutants compared with wild-type control. (C) Table showing the list of gene mutations in *(B)* identified with different color.

### TCS exposure regulates gene expression levels

In parallel, molecular mechanisms responsible for TCS-induced antibiotic resistance were determined by transcriptional analysis. Whole genome Illumina RNA sequencing was conducted on three types of mutants (*n* = 9) and wild-type *E. coli* (*n* = 3) in response to 0.2 mg/L TCS exposure for 8 h. Compared with non-treated wild-type *E. coli* (*n* = 3), acute exposure to TCS (*t* = 8 h) lead to representative changes in transcription among mutants and wild-type *E. coli* (Figure S1A and B). For wild-type *E. coli,* acute TCS exposure enhanced the expression of SIM network genes, such as *umuC* and *yfjY* that encode for DNA repair, efflux system-coding genes *acrE*, *mdtE*, *acrF*, *mdtB*, *mdtC* and *yddA*, and metallo-beta-lactamase superfamily coding gene *yjcS* (*p* < 0.05, Figure 3A, Tables S5, S6 and S7). In contrast, the expression of *ygiW*, *soxS* and *yhcN* genes that encode cellular antioxidants as well as membrane porin-encoding gene *ompX* were down-regulated (Figure 3A and Table S6 S7 and S8). These results suggest that at 0.2 mg/L TCS applies oxidative stress to *E. coli* while decreasing the expression of cellular antioxidant-encoding genes, which could then trigger the SIM response to DNA damage.

**Figure 3.**
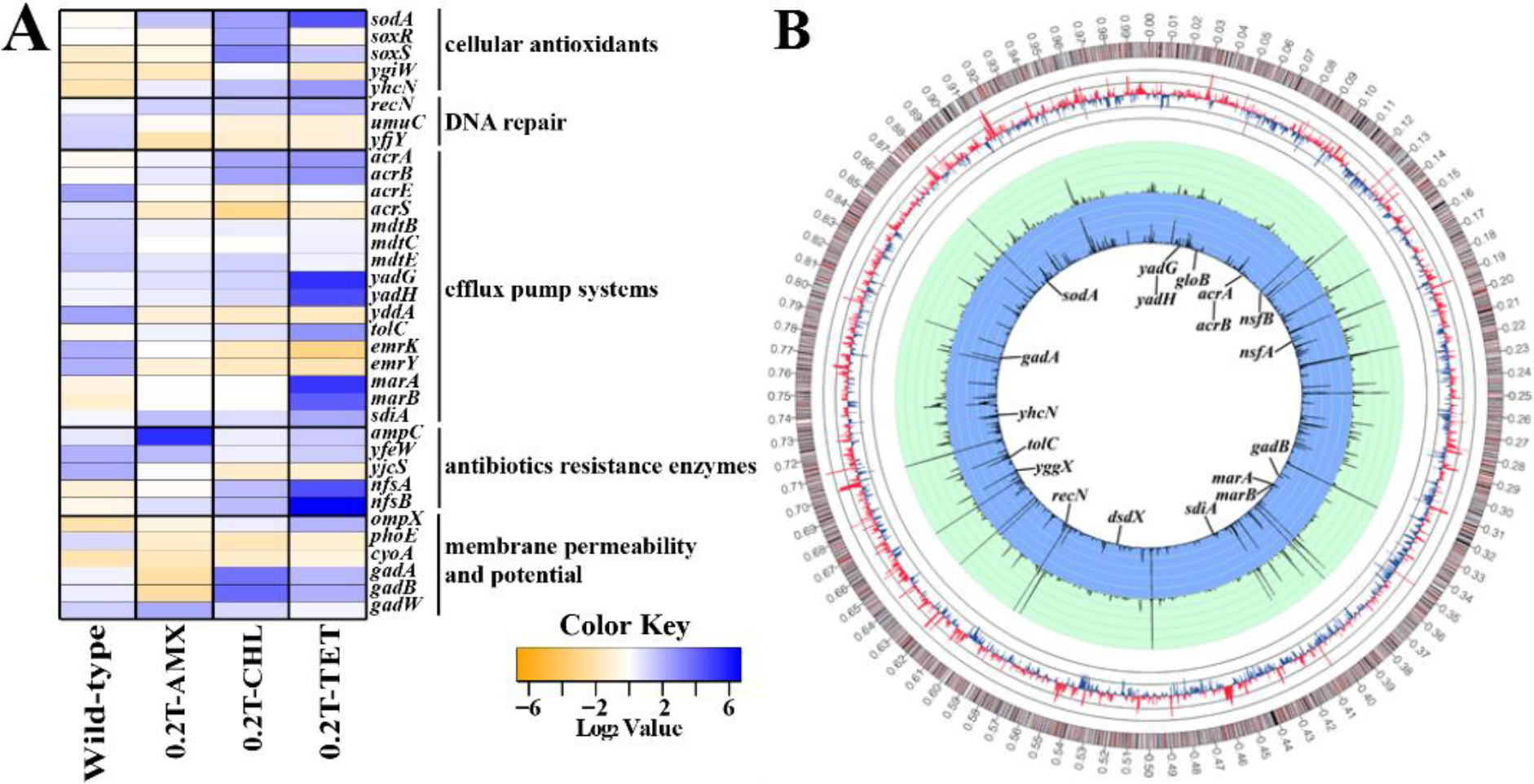
Gene expression with TCS exposure. (A) Differential expression of selected genes among wild-type (*n* = 1) and TCS-induced mutants (*n* = 8) which are influenced by TCS exposure. Differences in the levels of expression of stress-response and antibiotic resistance-related genes measured under control and 0.2 mg/L (*t* = 8h) TCS conditions are presented. Each gene is represented by the log_2_ of the fold change (i.e., RPKM TCS treated samples/untreated control ratio values). Orange represents the down-regulation of gene expression (log_2_ < −2), while blue represents up-regulation of mRNA expression (log_2_ > 2). (B) Global analysis of transcript levels in wild-type *E. coli* (*n* = 3) and 0.2T-TET mutants (*n* = 2) by RNA-seq. From inside out, the blue circle corresponds to the expression of each gene (represented as RPKM values) in 0.2T-TET mutants exposed to TCS-0.2 mg/L, while the green circle corresponds to wild-type *E. coli* grown without TCS. In both of the colored circles, thin circular lines represent an RPKM value of 1,000, with a limit at 5,000. The third line from the inside in the five gray circles represents the expression baseline of non-treated wild-type *E. coli*, and each thin gray circular line represent 4-fold changes (or log_2_ = 2) in expression of each gene. The outermost circle represents the full strain *E. coli* K-12 genome.

In TCS-induced mutants, genes encoding cellular antioxidants significantly increased (*p* < 0.05) expression levels when exposed to TCS, compared to non TCS-treated wild-type (e.g., *sodA* and *yggX*, Table S8), and this may lead to reduced ROS generation compared to the untreated *E. coli* (Figure 1I). In contrast, the expression levels of SIM-network gene *argD* and membrane porin-encoding gene *phoE* were significantly decreased when treated with 0.2 mg/L TCS (Figure 3A, Tables S5 and S7).

### TCS-induced mutations alter transcriptional response

TCS-induced genetic mutations may have caused increased antibiotic tolerance via regulating the expression of genes involved in antibiotic resistance (Figure 3A, Table S4 and S9). To illustrate, because the promoter of beta-lactamase-encoding gene *ampC* in *E. coli* K-12 overlaps the *frdD* gene region,^22^ the *frdD* mutation detected in 0.2T-AMX mutants may have affected the *ampC* promoter strength,^23^ thereby increasing the expression of *ampC* (log2 fold change (LFC) = 5.4), leading to the increased tolerance to beta-lactam antibiotics^24^ (Figure 1C, 2C, 3A and Table S7). In 0.2T-CHL strains, mutation in the *acrR* gene might have caused impaired function of *acrAB* repressor AcrR,^25^ resulting in increased transcription of the AcrAB multidrug efflux pump-coding genes *acrAB*.^26^ Additionally, mutation in the *soxR* gene may have increased the expression of *soxS,*^27^ resulting in an increase in efflux by up-regulating the *acrAB.*^28^ Therefore, overexpressed AcrAB multidrug efflux pump is likely to contribute to multiple antibiotic resistance^29^ (Figure 1C, 2C and 3A). In 0.2T-TET mutants, mutation in the *marR* gene may have attenuated binding efficacy of the *marRAB* operon repressor MarR,^25, 30^ leading to the overexpression of *marAB* genes which regulate global multidrug resistance.^31^ Consequently, MarAB may have initiated the expression of a cascade of antibiotic-resistance genes such as *acrAB*, *tolC*, *gadAB* and *yadGH,*^32^ which may have increased antibiotic resistance by promoting the expression and strength of the AcrAB-TolC efflux pump system (Figure 1C, 2C, 3A and B, Table S9).

### Possible mechanisms of antibiotic resistance induced by TCS exposure

In summary, our results suggest that at sub-MIC, TCS confers multiple antibiotic resistance in *E. coli* via ROS-induced mutation (Figure 4). Exposure to sub-MIC TCS (e.g. 0.2 mg/L) likely causes acute ROS generation in *E. coli*, and this oxidative stress triggers stress-induced mutagenesis that causes mutations in key genes such as *frdD*, *marR*, *acrR* and *soxR*. Those mutations result in either the overexpression of adjoin genes (*ampC* in the case of *frdD* mutation and *soxS* in the case of *soxR* mutation), or the enhanced expression of repressed genes (*marAB* and *acrAB*) caused by the mutations in repressor genes (*marR* and *acrR)*. Consequently, the enhanced expressions of beta-lactamase-coding gene *ampC* as well as local and global multidrug resistance regulator genes *acrAB*, *soxS* and *marAB*, may initiate the translation of beta-lactamase AmpC and multidrug efflux pumps AcrAB-TolC. Together with the reduced expression of porin-encoding gene *phoE* that affects cell membrane permeability, TCS-induced mutants could express extraordinary heritable resistance to a broad range of antibiotics by decreasing antibiotics uptake and increasing antibiotics efflux plus antibiotic degradation.

**Figure 4.**
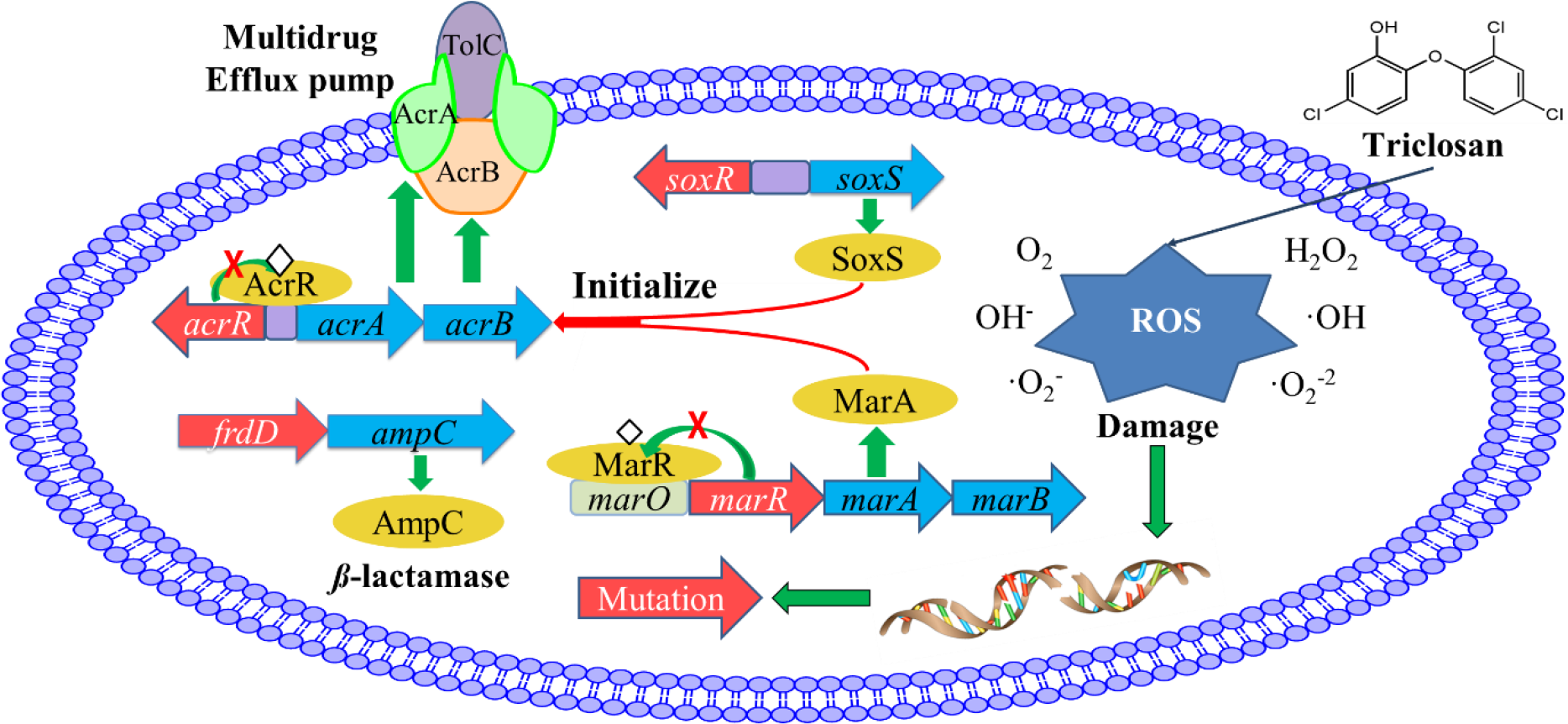
Proposed mechanism of TCS-induced antibiotic resistance. Oxidative stress generated by TCS can induce genetic mutations. Consequently, the genetic mutations lead to the overexpression of beta-lactamase-coding gene or global and local multidrug regulator genes that initiate the expression of multidrug efflux pumps. Red arrow represents gene mutations, blue arrow indicates up-regulated gene, yellow ellipse represents a protein, and white diamond means a loss of protein function.

Our findings suggest that non-antibiotic, antimicrobial chemical triclosan, at an environmentally relevant concentration, can induce multi-drug resistance with high hereditary stability. Considering the wide use of NAAM chemicals, and the prevalence of antibiotic-resistant bacteria, our study creates an imperative for a comprehensive analysis of the potential role of NAAM chemicals in triggering antibiotic resistance in microorganisms. To holistically evaluate the potential impact of NAAM chemical use on public health, a cohesive and rigorous understanding of the relationship between NAAM chemicals and global antibiotic resistance dissemination is critical.

## Notes

The authors declare no competing financial interest.

## ACKNOWLEDGMENTS

We acknowledge the Australian Research Council for funding support through Future Fellowship (FT170100196). Jianhua Guo would like to thank the support by UQ Foundation Research Excellence Awards. We are grateful to Dr. Eloise Larsen for reviewing and editing the manuscript. We thank Dr. Michael Nefedov of The University of Queensland for assistance with the BD FACSAria™ II flow cytometer and data analysis. This work was performed in part at the Queensland node of the Australian National Fabrication Facility.

## ABBREVIATIONS

NAAM: non-antibiotic, anti-microbial
TCS: triclosan
MIC: minimum inhibition concentration
ROS: Reactive Oxidative Species
SIM: stress-induced mutagenesis
AMX: amoxicillin
LEX: cephalexin
TET: tetracycline
CHL: chloramphenicol
LVX: levofloxacin
NOR: norfloxacin
KAN: Kanamycin
AMP: ampicillin
DCF-DA: 2’,7’-dichlorofluorescein diacetate
TBHP: N,N-dialkylanilines with tert-butyl hydroperoxide
SNP: single nucleotide polymorphism
FDR: false discovery rate
FPKM: fragments per kilobase of a gene per million mapped reads
LFC: log_2_ fold change

## Table of Contents

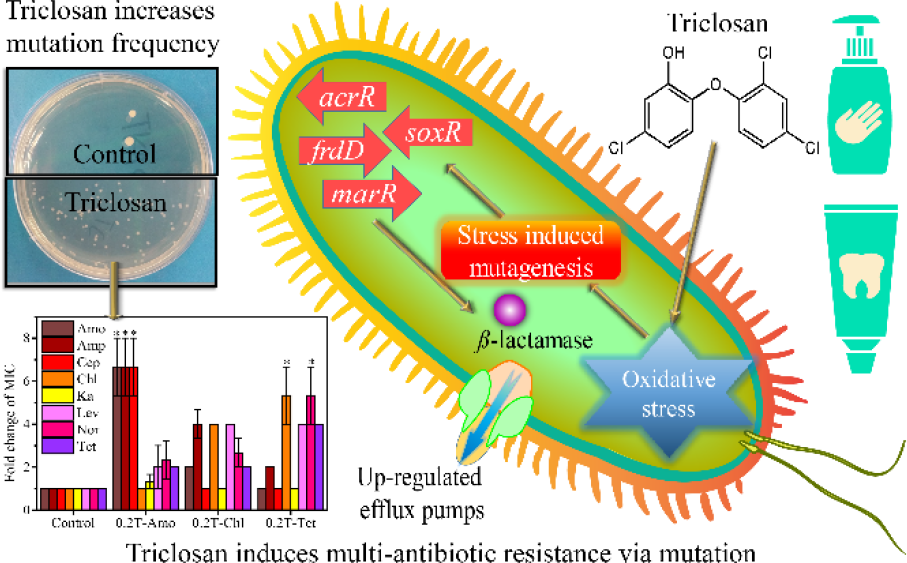

